# Evolutionary relatedness is a partially reliable predictor of host-associated microbiome composition

**DOI:** 10.64898/2026.09.08.749144

**Authors:** Kyle Buffin, Zakee Sabree

**Affiliations:** Department of Microbiology, The Ohio State University, Columbus, Ohio, USA; Department of Biological Sciences, University of Toronto Scarborough Campus, Toronto, Ontario, Canada; Department of Ecology and Evolutionary Biology, University of Toronto, Toronto, Ontario, Canada

**Keywords:** phylosymbiosis, insects, host-microbe interactions, coevolution

## Abstract

Multicellular animals emerged into a microbial world and they continue to be influenced by sympatric bacterial species. Host-associated microbiomes are recognized as critical contributors to their living domicile’s success. Detailing how microbiomes are assembled and maintained by their hosts has important implications for animal health, agricultural productivity, and biodiversity interventions across the tree of life. 10s-100s of bacterial genera are frequently detected in animal microbiomes, with some being uniquely and consistently detected in specific host species. Such discoveries have informed theories like phylosymbiosis that assert that host-associated microbiomes often reflect host species relatedness. We explored the predictive limits of this theory by performing a multi-scale dissimilarity-based comparative analyses using publicly-available microbiomes drawn from 113 insect species spanning 14 insect orders. We detected a moderate inter-order phylosymbiotic signal (Mantel test: r = 0.392, p<0.001) that was weakened (r = 0.173, p<0.001) when the Blattodea were excluded from the analysis. Intra-order level phylosymbiotic signals were detected for lepidopteran, hymenopteran, hemipteran, and blattodean host species microbiomes. These signals were often explained by obligate endosymbionts and/or nestmate-acquired gut bacteria, rather than by the total microbiomes. Host species-defining bacterial genera often represented a fraction of the total observed microbial diversity in their respective insect host species. We hypothesize that members of insect orders can exhibit considerable microbiome compositional variability where host speciation events may be minimally reflected in their microbiomes taxonomic membership, but host species-defining bacterial genera are likely when host behaviors or physiologies that facilitate reliable intra- and inter-generational bacterial transmission or acquisition are present.

**IMPORTANCE:** While all animals have microbiomes, the principles determining their composition remain an incomplete story. We leveraged the diversity of Insecta, arisen from 400 My of evolution, to evaluate if microbiome evolution is synchronized with the evolutionary history of their associated hosts. We found that host evolution plays a limited role in predicting microbiome composition, particularly due to non-uniformity of host-microbial symbioses across insects. While Blattodea in particular harbor exceptionally diverse microbiomes, many insect guts are characterized by few bacterial genera. Our results suggest that host evolutionary history is but one of many factors, including differences in microbiome transmission modality, influencing insect microbiome evolution.

## INTRODUCTION

The importance of relationships between host organisms and their associated microbial communities cannot be understated, and detailing the mechanisms driving their composition and function clarifies important underpinnings of these host-microbial symbioses. Metazoans emerged into a world shaped by their bacterial predecessors, and they have evolved structures and function that facilitate cohabitation, collaboration and competition with sympatric microbes. Host-associated bacterial communities are assumed to be assembled either stochastically or through mechanisms evolved over the history of the species and fixed within the host’s genetics [1–3] [4, 5]. Phylosymbiosis is a hypothesis that specifies that host phylogenetic relatedness and microbiome community relatedness are positively correlated, which implies that closely related host species tend to harbor more similar microbial communities [2, 6]. Given its intuitive charm, the utility of this hypothesis in explaining host-associated microbiome composition has been explored across metazoan species that include mammals [7–10], birds [7, 11], insects [12, 13], invertebrates [14, 15], and plants [16, 17]. Phylosymbiosis at its core implies that the mechanisms structuring host-associated bacterial communities are, at least in part, conserved within closely-related host species, yet this hypothesis does not prescribe specific mechanisms. Several modalities for host-associated microbiome acquisition have been detailed in different host systems, which includes parental intergeneration transmission, filial intrapopulation transmission and environmental uptake-plus-host filtering [18, 19]. Microbes being systematically passed from one generation to the next, also termed ‘vertical transmission,’ could result in a pattern of microbiome compositions across sister host species predicted by phylosymbiosis [3, 6]. In host species lacking evidence of vertical bacterial transmission, phylosymbiosis may still be observed when the hosts employ molecular (e.g. immune system), chemical (e.g. pH) and mechanical (e.g. proventriculus) mechanisms that filter and select for specific gut microbiota [18]. Additionally, similarity in microbiome composition between similar hosts may also be driven by shared ecology, such as dietary and behavioral traits which restrict each host’s exposure to microbial diversity [20, 21]. Microbiome compositions across closely-related host species that do not fit a phylosymbiotic pattern of similarity may suggest that factors like dietary shifts, habitat transitions, or horizontal microbial acquisition are disrupting phylogenetically conserved host-microbiome relationships.

Insects are the most species-rich class of extant terrestrial animals and inhabit an incredible diversity of ecological niches. Many insects foster close associations with bacteria that are essential for their development and survival [22]. Insect gut microbiomes are marked by compositional differences suggestive of varying levels of host specificity. Honey bees, bumble bees, and aphids for example, have highly consistent microbiomes comprised of tens of bacterial species [23, 24], while cockroaches and dung beetles have been observed to have microbiomes with hundreds of bacterial species [25, 26].

Several insect species domicile bacteria (e.g. *Buchnera aphidicola* [27], *Blochmannia* [28], and *Wigglesworthia glossinidia* [29]) that are restricted to specialized tissues or cells within their hosts, and these insect hosts rely on many of these bacteria for normal growth and development. These endosymbionts have highly reduced genomes, are strictly vertically transmitted and are integrated into their host’s metabolism – all of which reflect millions of years of coevolution [30]. Extracellular vertical transmission [31], coprophagy [32], environmental uptake-plus-filtering [33] are among other modalities of microbiome assembly observed in insects [34]. Given their ecological diversity, the complexity of their interactions with microbes, and some well-known microbiome transmission modalities, insects provide an excellent case study for examining the degree to which known host species relationships are reflected in their microbiomes. Drawn from publicly-available insect-associated microbiomes, we investigated the efficacy of host evolutionary history for predicting gut microbial community compositions over nearly 400 My of insect evolution.

## MATERIALS AND METHODS

A comprehensive meta-analysis of publicly available 16S rRNA amplicon sequencing data was conducted to characterize insect-associated bacterial communities across the broadest possible phylogenetic range of host species. Sequencing projects were identified through manual searches of the NCBI Sequence Read Archive and curated to retain only those targeting internal microbial communities using Illumina sequencing platforms. Projects were excluded when samples could not be reliably attributed to internal tissues, untreated adult specimens, or when insufficient metadata prevented meaningful classification. A full list of included BioProjects is provided in **Supplemental Materials**.

Sequencing data were obtained using the SRA Toolkit (v3.0.0) and processed independently per BioProject within the QIIME 2 framework (v2024.10) [35]. Primer sequences were identified from a curated library of common 16S primers and trimmed with Cutadapt [36]. Quality filtering was applied using a minimum q-score threshold of 15 with a 4 bp sliding window. Amplicon sequence variants (ASVs) were resolved using DADA2 [37], and taxonomy was assigned using a pre-trained Naive Bayes classifier against the SILVA 138 99% database [38]. Non-bacterial sequences including eukaryotic, chloroplast, and mitochondrial assignments were removed, as were ASVs with fewer than 11 total reads across all samples, and samples with fewer than 10,000 reads. Merged ASV tables were collapsed to genus level to facilitate cross-project comparisons. Only samples of adult specimens were retained, and host species represented by fewer than 10 samples were excluded from downstream analyses.

To distinguish consistently host-associated bacteria from transient taxa, we developed a bootstrap-based prevalence framework. For each host species, 1,000 bootstrap iterations were performed by resampling available samples with replacement; the proportion of samples in which each bacterial genus was detected was recorded per iteration. A taxon was classified as prevalent for a given host if the value 0.80 fell within the 95% bootstrapped confidence interval of its prevalence, indicating a consistent and reliable association. This procedure was additionally applied to subsets of samples stratified by tissue, sex, habitat, life stage, and source BioProject when sample sizes permitted (n ≥ 10 per stratum). Full details are provided in **Supplemental Materials**. To better understand our identification of core bacteria the relative abundances of all core genera within each microbiome sample for host species were summed to estimate how much of the total community is comprised of core genera.

Phylosymbiosis was assessed using Mantel tests (999 permutations, vegan package in R [39]) to correlate host cophenetic phylogenetic distance derived from a TimeTree-generated [40] host phylogeny against pairwise microbiome dissimilarity. Dissimilarity between host species was quantified as pairwise distances between ordination centroids for each host’s sample cloud, calculated separately for three community partitions (whole, core, and non-core communities) under both abundance-weighted and binary (presence/absence) frameworks, yielding six total comparisons. Partial Mantel tests controlling for dietary dissimilarity, based on Gower distance, were used to evaluate the contribution of phylogenetically conserved diet to observed phylosymbiosis. Robustness was assessed via jackknife analysis by iteratively excluding each insect order and recalculating Mantel test statistics. Order-level Mantel tests for orders represented by at least five host species was also determined. Complete details of all statistical procedures are provided in **Supplemental Materials**.

## RESULTS

### Dataset overview and defining core genera

Microbiomes from 113 insect species, including *Daphnia magna* as an outgroup, spanning 14 insect orders were analyzed **(Fig. 1A, Dataset S1)**. Each insect host species was represented by at least ten, and up to 564, bacterial community samples comprised of at least 10,000 high-quality 16S rDNA amplicon reads derived from nucleic acids extracted from adult specimens **(Supplemental Materials)**. Amplicon sequence variants assembled from high-quality reads were taxonomically-assigned using Arb-SILVA (version 138), and genus-level phylotypes (hereafter referred to as ‘phylotypes’) were used to describe taxa comprising the host-associated communities to enable comparisons across diverse sequencing projects.

**Figure 1:**
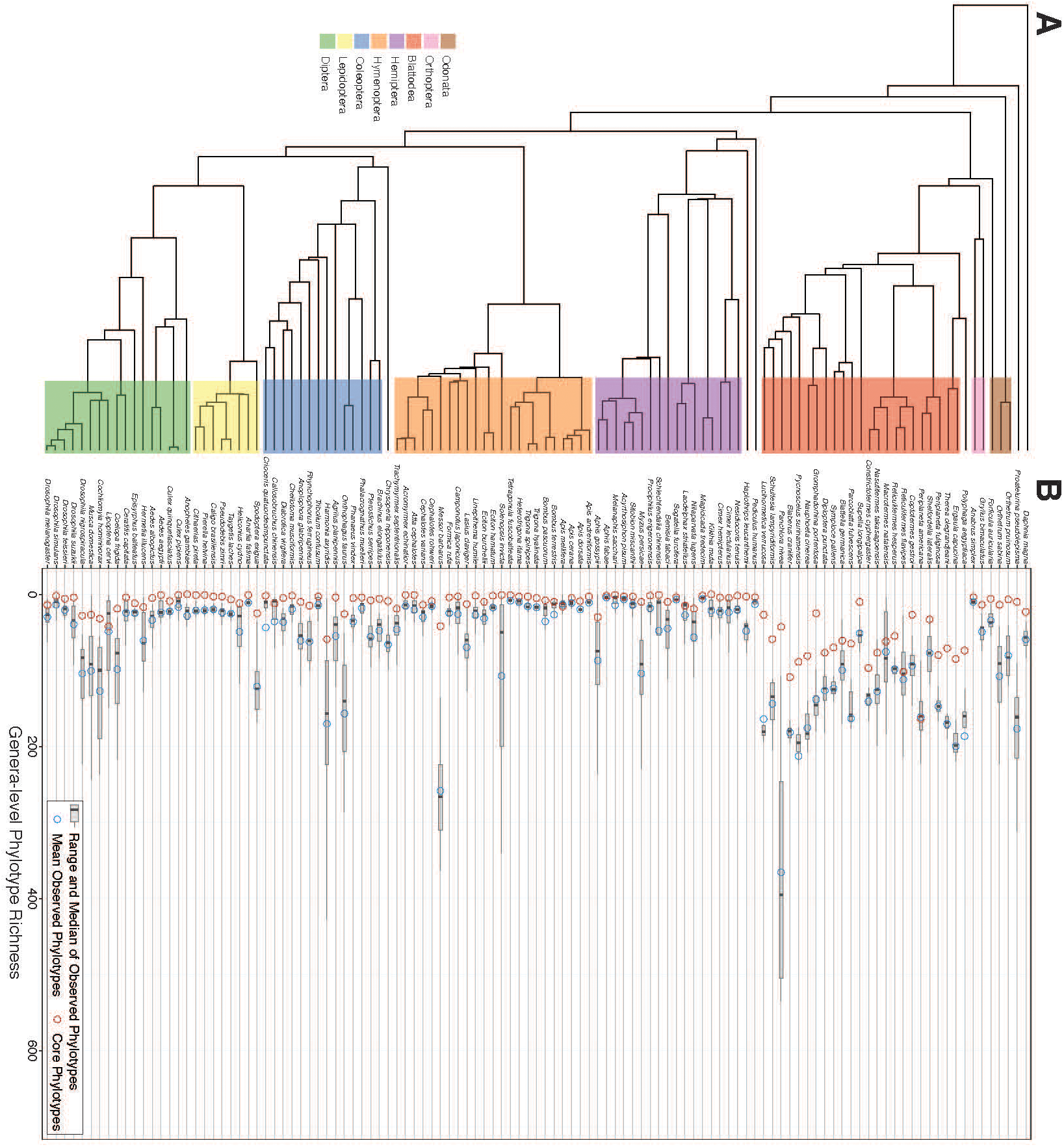
Diversity of host-associated bacterial communities varies widely across insect phylogeny. Our dataset is comprised of 113 host species represented by at least ten bacterial community samples. These host species represent 14 different taxonomic orders. Host order when inclusive of more than one species is indicated on the host phylogeny by the colored bar. The number of core phylotypes (red circles) determined for each host species were consistently less than the mean (blue circles) and median (box plot) number of observed phylotypes.

Phylotype richness (i.e. total observed genera) per host species indicated that alpha diversity spanned nearly three orders of magnitude, with termites and cockroaches (Insect order: Blattodea) exhibiting the highest average phylotype richness (range: 51.5-365.5) **(Dataset S1)**. We defined ‘core’ phylotypes by their detection in ≥80% of samples per gut microbiome sequencing project per host species, irrespective of their relative abundances in those samples (note: any phylotypes represented by <10 reads in a sample were removed from those samples upstream; **see Supplemental Materials** for methodological details). Core phylotype richness was consistently less than the average total phylotype richness for each host species **(Fig.1B)**, with Blattodean host species harboring the most core phylotypes (average: 93.85, range: 10-162) while >50% of remaining host insect species had fewer than ten core phylotypes **(Dataset S1)**. Collector’s curves of core phylotypes indicated that biases due to differences in number of sample replicates per host were minimal at the minimum sampling depth per species (n=10) **(Fig. S1A-E).** The average number of observed phylotypes roughly predicted the number of core phylotypes for a given host **(Fig. S2**); however, this predictability diminished as phylotype richness increased. As observed phylotype richness increases, core phylotype richness becomes more variable. While Diptera and Coleoptera exhibited per species phylotype richness of 48.2 and 58.1, respectively, and order-level average core phylotype richness of 13.1 and 11.4, respectively, many host species within these orders exhibited considerable differences between average observed phylotype richness and core phylotype richness **(Fig. 1)**.

### A few bacterial phyla represent most core microbiota

Core phylotypes represented ≥50% of amplicons in microbiome samples for 88 out of 113 host species **(Fig. 2A)**, which included most-to-all host species assigned to Blattodea, Hemiptera, Hymenoptera, Odonata, and Orthoptera orders. In contrast, the total relative abundances of core phylotypes varied widely per microbiome sample for lepidopteran, dipteran and coleopteran host species, with both multimodal and uniform distributions being observed **(Fig. 2A)**. When present, obligate endosymbiotic bacterial genera (e.g. *Buchnera, Sulcia, etc.*; **Dataset S2**) were abundant (i.e. >0.5% of amplicons per sample) and amongst, if not the sole, core phylotypes within their host species’ microbiomes. In contrast, obligate endosymbionts in turtle ants and carpenter ants (*Blochmmania*) and cockroaches (*Blattabacterium*) were amongst the 10s-100s of core phylotypes, representing a relatively small fraction of the total reads per host species. While some endosymbiont-bearing insect species maintain low-diversity microbiomes with one-to-few endosymbionts, others commonly domicile a diverse gut microbiota despite having endosymbiotic mutualists.

**Figure 2:**
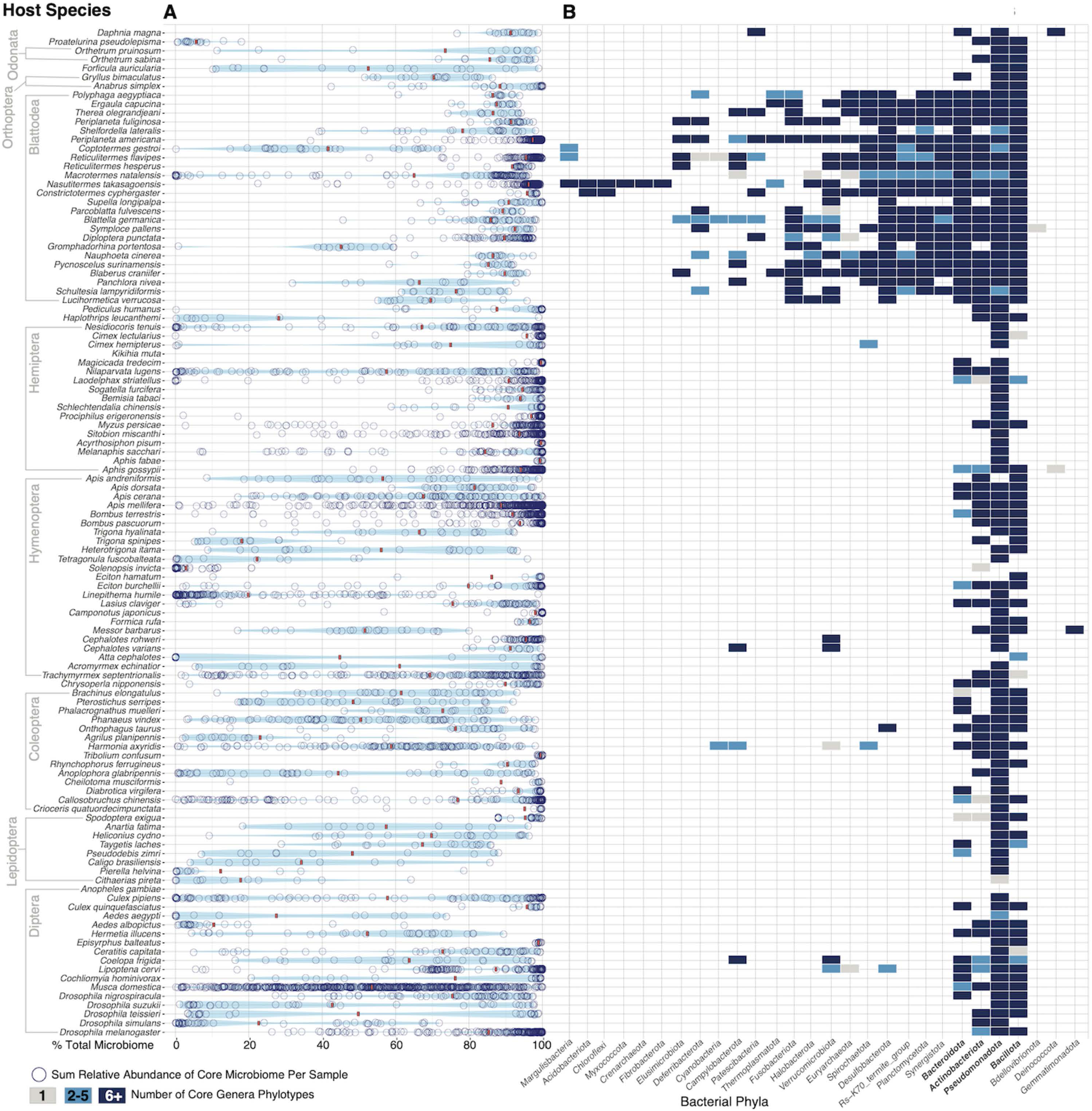
Per sample abundance and taxonomic distribution of core microbiota across insect hosts. (A) Core phylotypes were determined per host species as described and the sum relative abundance of all host-specific core microbiota per sample was determined (Blue circles). Mean relative abundance of core phylotypes per host species is indicated by red bar. (B) Distribution of host-specific core microbiota across taxonomic phyla. Intra-phylum diversity is shown by box color. Bolded bacterial phyla (Pseudomonadota, Bacteroidota, Bacillota, Actinoycetota) represent those most conserved across the entire insect phylogeny.

510 unique core phylotypes were detected across 113 host species, with most being assigned to the Pseudomonadota, Bacteroidota, Bacillota, and Actinobacteriota phyla **(Fig. 2B).** Pseudomonadota genera were the most widespread, with at least one representative core phylotype in 106/113 host species **(Fig. 2B)**. Blattodean insects uniquely harbored core phylotypes from up to 14 additional bacterial phyla, with core phylotype richness for shared phyla (i.e. Pseudomonadota and Bacillota) additionally being the highest in blattodean host species **(Fig. 2B)**. Despite Pseudomonadota, Bacteroidota, Bacillota, and Actinobacteriota-associated core phylotypes being detected in host insects, 42.5% of all core phylotypes were uniquely detected in a single host species **(Fig. S2).**

### Core phylotypes are driving phylosymbiosis across the insect phylogeny

Six datasets describing host associated bacterial community composition were built to evaluate pan-insect host-microbiome phylosymbiosis: abundance-weighted and binary-transformed; all genus-level phylotypes (‘total’ community), core phylotypes (‘core’) and non-core (‘core’ subtracted from ‘whole’) phylotypes. Across these six datasets, dissimilarity was determined by finding the centroid of all microbiome samples for a given host, before measuring the pairwise distance to each other host’s respective centroid. When using abundance-weighted datasets, the whole community exhibited the strongest phylosymbiotic signal (total: r=0.312, p<0.001, core: r=0.235, p<0.001, non-core: r=0.061, p=0.055) **(Fig. 3A).** Repeating these analyses using binary-transformed datasets also resulted in the total community having the strongest signal, with the core dataset having a similar test statistic (total: r=0.392, p<0.001, core: r=0.365, p<0.001, non-core: r=0.117, p=0.006) **(Fig. 3B)**.

**Figure 3:**
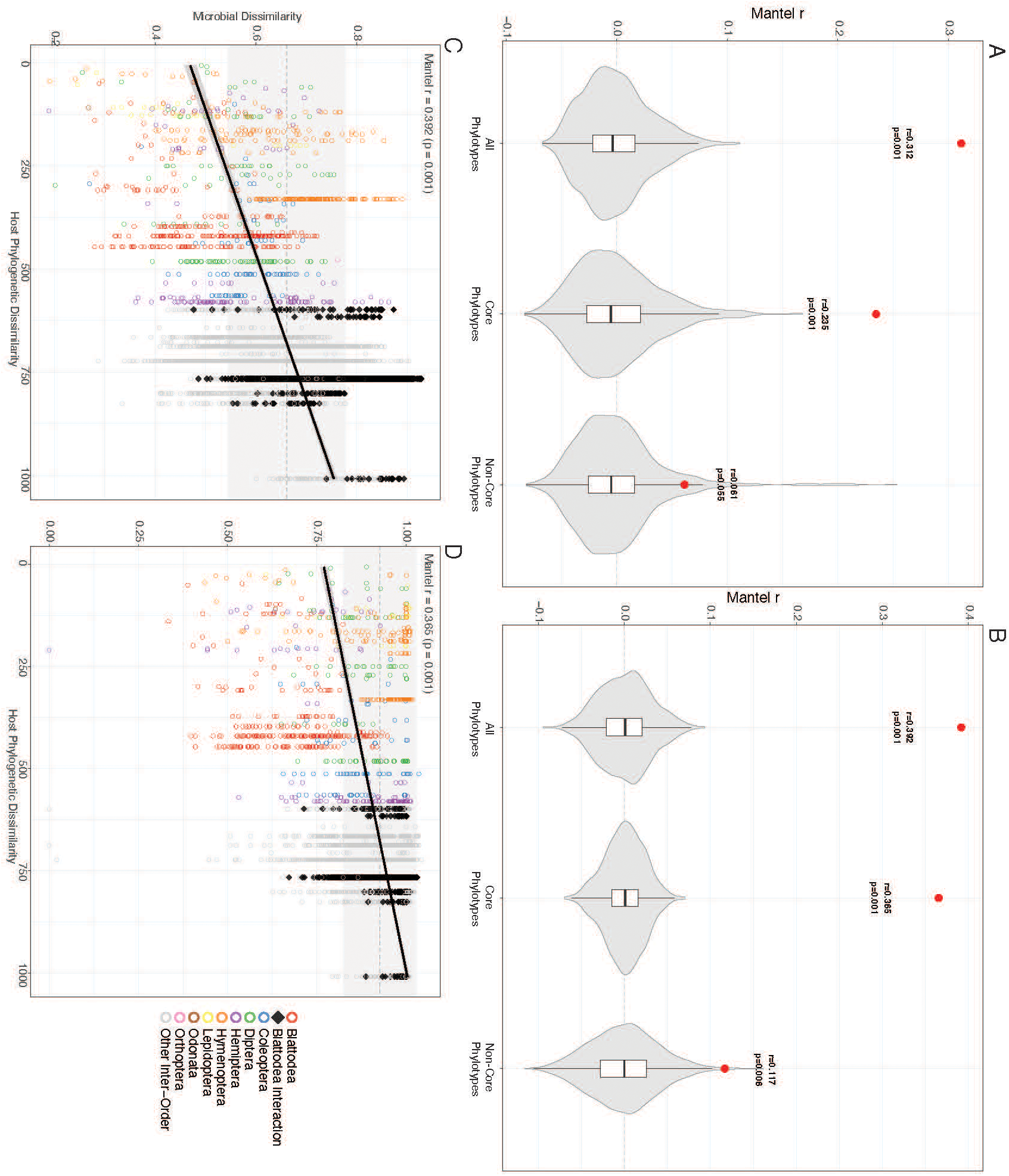
Determination of phylosymbiosis across insects using different microbial dissimilarity models. **A and B** Comparison of phylosymbiosis measured between three datasets; 1. Whole community, 2. Core community and 3. Non-core community. Microbiome dissimilarity was calculated between species by taking the pairwise distances of each host’s ordinational centroid for either (A) abundance weighted Aitchison distance or (B) Jaccard dissimilarity. For both methods the whole community (left) dataset resulted in the highest level of measured phylosymbiosis using mantel tests. **C and D** Phylosymbiosis graphically represented as a correlation between host phylogenetic dissimilarity and microbiome dissimilarity for both whole (C) and core (D) communities. Overall, the core community dissimilarity is higher between all host species (pairwise values closer to 1). The points are colored to represent pairwise comparisons between two host species within the same order, or comparisons between Blattodea and any other order (Black).

Hierarchical clustering applied to dissimilarity matrices constructed from the total, core and non-core binary-transformed datasets yielded dendrograms that illustrated the degree to which insect host microbiomes recapitulated known host species phylogenetic relationships **(Fig. 4)**. Total and core microbiome-based dissimilarity matrices yielded dendrograms in which the Blattodea order was recovered, with some internal rearrangements and recruitment of *Gryllus bimaculatus* crickets. With the exception of *Spodoptera exigua,* the Lepidoptera order was primarily recovered with the total microbiome **(Fig. 4)**. Bee species relationships were preserved in the total microbiome-based dendrogram, and partially in the ‘core’ dendrogram. Ant species were erroneously dispersed across the dendrogram rather than in proximity to other hymenopterans, demonstrating poor recovery of host species relationships using any of the microbiome datasets. Amongst other orders some congeneric groups were recovered (e.g. *Cimex* and *Orthetrum*), depending on the dataset used, but examples to the contrary (e.g. *Drosophila, Aedes, Culex*) were also observed.

**Figure 4:**
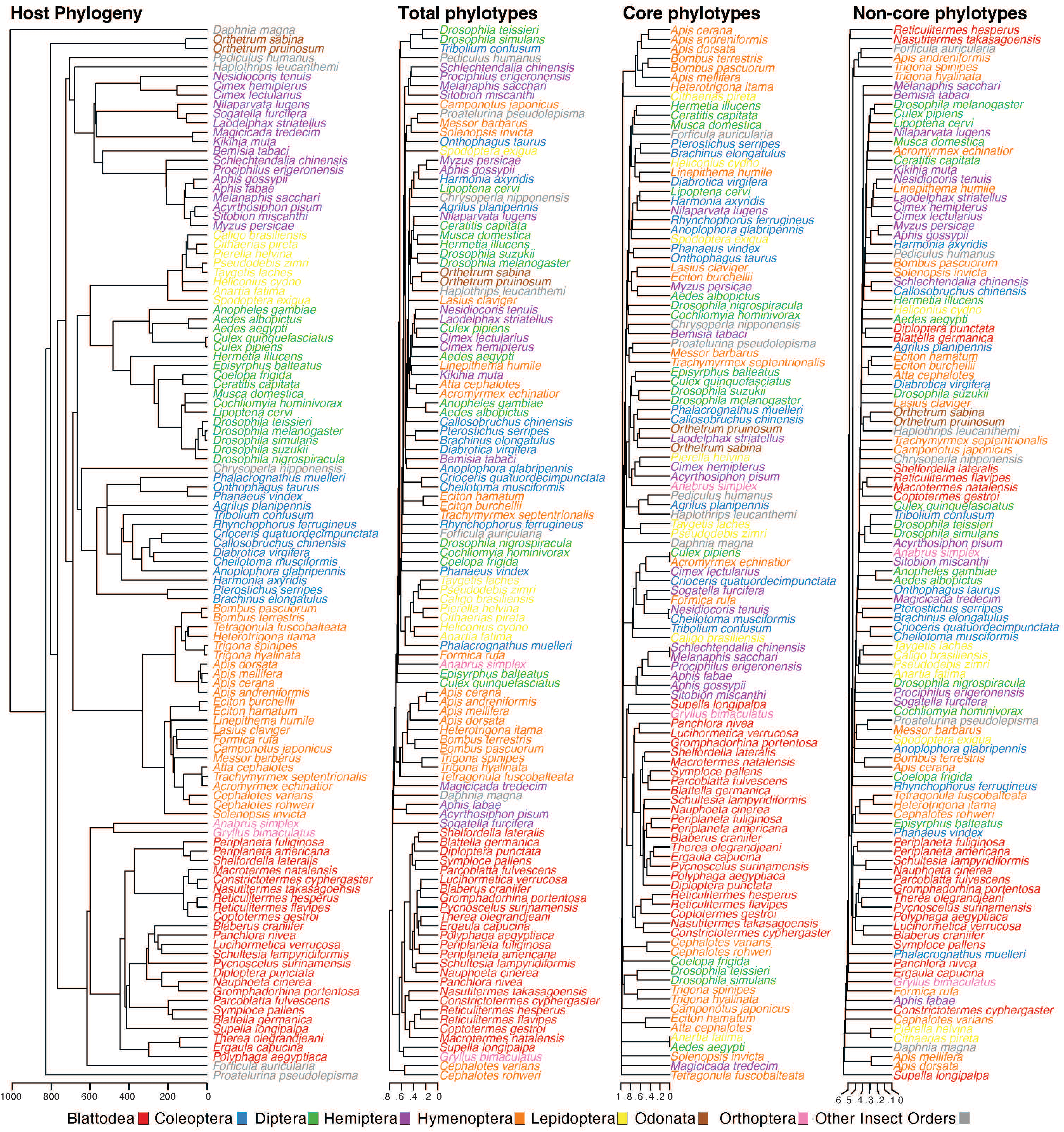
Insect bacterial community compositions largely do not recapitulate host phylogenetic relationships. Hierarchical clustering was applied to the dissimilarities between host species bacterial communities for the whole, core, and non-core datasets and compared to the host phylogeny. Longer branch lengths indicate greater dissimilarity between hosts. Host order when represented by more than 1 species is shown with different colors. While some taxonomic groups of insects cluster strongly across all datasets (Blattodea, Apidae), there are generally limited recapitulations of host relationships across all insects.

### Blattodean microbiomes defined pan-Insecta phylosymbiosis

Blattodean microbiomes are distinct from those of other insect hosts **(Fig. 2)**, suggesting that they drive the observed pan-insect phylosymbiotic signal. A leave-one-out resampling approach was used to evaluate this prediction where single insect orders were removed from the datasets and the measured phylosymbiosis was re-assessed. Removal of blattodean microbiomes from the dataset eliminated most of the phylosymbiotic signal across all microbiome datasets, while the removal of other insect orders had zero-to-less than the same magnitude of effect **(Fig. 5A)**. The baseline whole community r-statistic of 0.392 decreased to 0.173 after the removal of Blattodean microbiomes. As demonstrated, Blattodean bacterial communities are very different compositionally from those of every other insect species. This, and the phylogenetic dissimilarity between most of the included insects and blattodeans drove the phylosymbiotic signal (**Figs. 3C and 3D)**. Intra-order phylosymbiosis was measured to evaluate if a strong phylosymbiotic signal within Blattodea contributed to the effect of their removal, in addition to examining if there is evidence of phylosymbiosis across smaller groups of insects. Within-blattodean phylosymbiosis (all phylotypes: r=0.249, p=0.004; core phylotypes: r=0.191 p=0.20) was below the pan-insect baseline (all phylotypes: r=0.392, p<0.001, core phylotypes: r=0.365, p<0.001). However, relatively higher intra-order phylosymbiotic signals were observed for hymenopteran whole communities (all phylotypes: r=0.611, p<0.001), hemipteran core communities (core phylotypes: r=0.554, p<0.001) and lepidopteran whole and variable (all phylotypes: r=0.914, p=0.006, non-core phylotypes: r=0.543, p=0.006) microbiome groups **(Fig. 5B)**.

**Figure 5:**
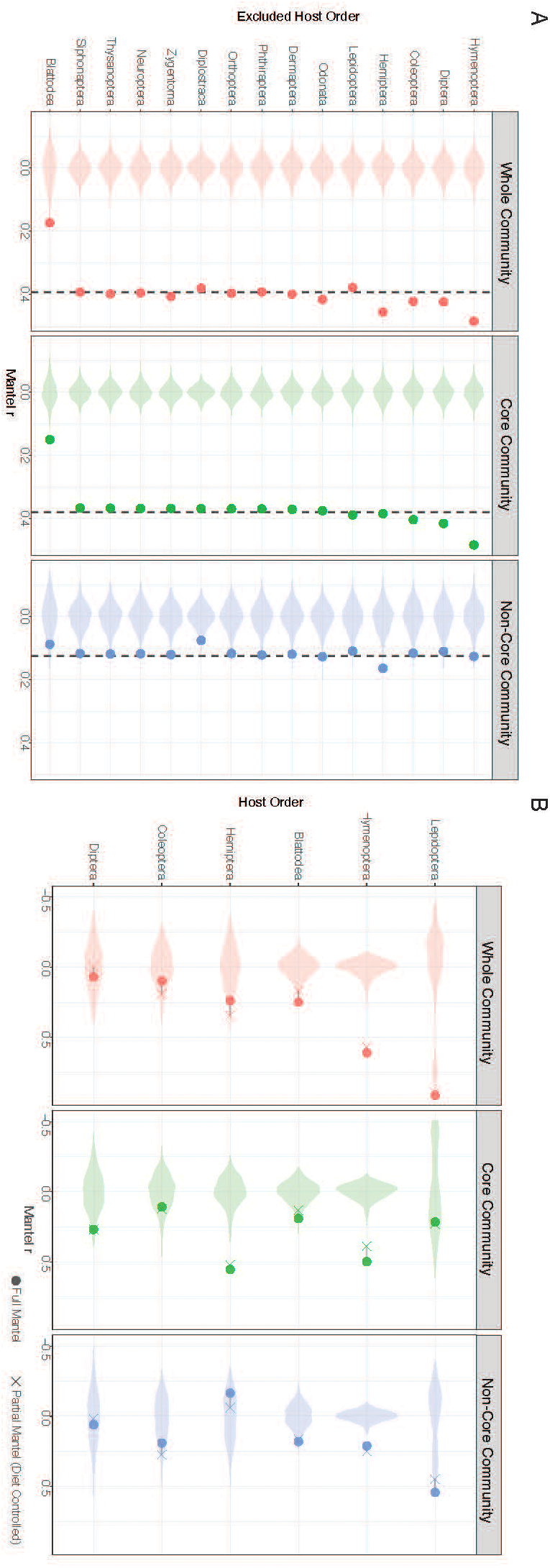
Inter-order differences are the primary driver of insect phylosymbiosis. **(A)** To evaluate the stability of insect phylosymbiosis and the contribution to phylosymbiosis by different insect orders, a jackknife approach was completed by consecutively removing each insect order from the dataset and recalculating phylosymbiosis. This was done across all three community datasets. In the whole community and prevalent community particularly, the removal of Blattodea hosts significantly decreases the measured phylosymbiosis relative to the dotted line representing the baseline measure across all insects. **(B)** Intra-order phylosymbiosis was also measured for each insect order with at least 5 host species. The measured phylosymbiosis across insects seems to be reliant on the inclusion of Blattodea in the dataset, however within Blattodea there is no evidence of phylosymbiosis. In fact, across most insect orders the within-order phylosymbiosis is non-existent or lower than what was observed across all insects. Hymenoptera and Hemiptera contrast these results, demonstrating increased intra-order phylosymbiosis in some datasets.

### Dietary similarity partially explains insect phylosymbiosis

The average richness of total and core phylotypes in diet-categorized host species revealed both intra- and inter-diet variations, with reduced variation in the core-only phylotype averages **(Fig. S3)**. Microbiomes of herbivorous insects, especially plant sap feeding insects, exhibited relatively low total and core phylotype richness, while xylophagus and omnivorous insects had relatively higher phylotype richnesses, with the latter insects exhibiting the greatest phylotype richness variation **(Fig. S3**). Xylophagous and omnivorous diet groups were largely comprised of blattodean insects, and the pattern of their uniquely high diversity communities can be seen in both the whole community and core phylotype diversity measures. Phylosymbiotic signals for abundance-weighted or binary-transformed whole community and core phylotype datasets were slightly decreased when host species’ diet was controlled for using partial mantel tests **(Fig. S3)**. For both abundance-weighted comparisons (based on Aitchison dissimilarity), no difference in Mantel’s r between the full and partial tests was observed **(Fig. S3)**. For the binary-transformed comparisons the partial Mantel test result was lower for both the whole community (full: r=0.392, partial: r=0.347) and core community (full: r=0.365, partial r=0.305). As such, dietary similarity appears to have a limited but significant impact on similarity of microbiome composition for closely related insect lineages. Within-diet group phylosymbiosis was observed in three generalized dietary categories; herbivores (whole: r=0.358, core: r=0.347), carnivores (whole: r=0.647, core: r=0.653), and omnivores (whole: r=0.677, core: r=0.676). Removal of blattodean species from these comparisons diminished the phylosymbiotic signal for both herbivorous whole and core communities (whole: r=0.225, core: r=0.290) as well as omnivorous core communities (whole: r=0.413, core: r=0.080) **(Fig. S3)**. While a strong intra-diet signal for carnivorous insects (whole: r=0.647, core: r=0.653) was observed, low species representation (n=9) across very diverse insect orders likely explains this result.

## DISCUSSION

### Cockroaches and termites consistently maintain exceptionally high gut bacterial diversity

Pan-insecta phylosymbiosis was primarily defined by the Blattodea in these analyses. Cockroach and termite microbiomes harbored, on average, an order of magnitude more core genera-level phylotypes than detected in all other insect orders, with many of these phylotypes being uniquely associated with blattodean species despite their hosts occupying distinct ecological niches (i.e. wood-feeding vs. omnivorous). Core phylotype richnesses of termite- and cockroach-associated microbiomes were similar, despite the former being primarily xylophagous. However, diet alone did not explain the relatively high core phylotype richness of blattodeans given that other omnivorous insect species included in this study (e.g. carpenter ants, dung beetles and crickets) harbored fewer core phylotypes despite some within-insect species microbiome samples exhibiting high total phylotype richness **(Fig. 1)**. Blattodean core phylotypes most often represented >80% of amplicons in microbiome samples **(Fig. 2).** While a similar trend was observed in many hemipterans and some bees, beetles and *Drosophila* **(Fig. 2)**, the microbiomes of these insect species were much less diverse and often dominated by vertically-transmitted endosymbionts and filially transmitted gut symbionts. We expect that the maintenance of highly diverse core microbiomes across blattodean species is a trait ancestral to the divergence of cockroaches and termites, which was estimated to be >200-250 M.y.a. [41], and this core phylotype richness has been maintained through several speciation events, resulting in hundreds of termite and cockroach genera [42]. Filial and parental coprophagy, proctodeal trophallaxis, and gregarious domiciling are host behaviors commonly exhibited by blattodeans that could enable moderate-to-high fidelity intra- and inter-generational gut microbiome transfer [43–47]. Additionally, bacteria-linked volatile organic compounds (VOCs) in cockroach feces that act as attractants, as observed in *Blattella germanica [48]*, could increase the likelihood of filial coprophagy and enhance microbiome compositional consistency over space and time. Several anaerobic genera, including those within the *Synergistota, Desulfobacterota*, *Fusobacteriota*, *Spirochaetota*, were uniquely detected as core genera in most-to-all blattodeans, which potentially reflects their reliance upon trophallaxis, coprophagy and gregarious domiciling to persist within their hosts. These and other core phylotypes distinguished the blattodean order **(Fig. 3C-D)**. Despite the uniqueness of blattodean microbiomes, we observed limited recapitulation of known blattodean host species relationships **(Fig, 4)** and thereby weak intraorder phylosymbiotic signal **(Fig, 5B)** indicating that transmission of bacterial diversity between generations may dominate transmission of host-specific species. While further investigation is needed to ascertain the origin of the unusually diverse gut microbiomes found within blattodean host species, we hypothesize that inter- and intra-generational microbiome transmission mechanisms, like coprophagy and trophallaxis, contribute to the maintenance of order-defining core genera in insect gut microbiomes.

### Host species evolutionary relationships are only moderately useful predictors of insect microbiome composition

Exclusion of the Blattodea from pan-Insecta phylosymbiosis analyses reduced the signal by nearly half (**Fig. 5A**), which suggested that microbiomes from many divergent insect orders in a single data set could not be used to reliably recapitulate all known host species relationships **(Fig. 4)**. In contrast, intraorder phylosymbiotic signals were relatively strong for some orders when using all phylotypes (e.g. lepidopterans and hymenopterans) or core phylotypes (hymenopterans and hemipterans) (**Fig. 5B**), which we attribute to the obligate endosymbionts (e.g. *Buchnera*, *Wolbachia*, *Sulcia*, *Hodgkinia, Liberibacter*, *Hamiltonella, Portiera*) and vertically- and/or community-acquired bacteria (*Bombilactobacillus, Lactobacillus, Gilliamella*, *Snodgrassella, Bombella,* Erysipelotrichaceae ZOR0006, *Verticiella, Parapusillimonas, Ventosimonas, Cephaloticoccus*) harbored by hemipterans and hymenopterans. The similarities between bee microbiomes and dissimilarities between ant microbiomes **(Fig. 4)** likely explains the intraorder phylosymbiotic signal observed for hymenopterans. Despite their relative similarities to each other, bee microbiomes are known for being highly dynamic and exhibiting symbiont gains and losses across and within bee genera (e.g. *A. mellifera*, *A. cerana*, *A. dorsata*, *A. florea* and A*. andreniformis*; [49]) which can complicate co-diversification claims at various taxonomic resolutions. In this study specifically, bee-associated bacterial compositions generally appear similar because of the genus-level phylotype being used to characterize them. Broadly, given that ants and bees include many eusocial lineages, these results further highlight how horizontal transmission of microbiomes through filial social behaviors, as in termites and cockroaches, could lead to compositional consistency [50].

Dipteran and coleopteran microbiomes poorly recapitulated known host species relationships within the pan-Insecta analysis **(Fig. 4)** and their intraorder phylosymbiotic signals were among the lowest observed **(Fig. 5B)**. In most cases the underlying host microbiome samples were frequently comprised of peripheral phylotypes that, in sum, could represent >50% of the amplicons per sample per dipteran or coleopteran host **(Fig. 2).** Members of the Pseudomonadota and Bacillota constituted the dominant low-abundance core phylotypes across dipterans (e.g. *Gluconobacter, Acetobacter, Lactobacillales*, *Asaia*, *Wolbachia, Acinetobacter and Pseudomonas*) and coleopterans (*Vagococcus, Wolbachia, Pseudomonas, Acinetobacter*). We expect that the low proportion of core phylotypes in coleopteran and dipteran microbiomes reduces their ability to recapitulate their host’s evolutionary histories **(Fig. 4).** Although fecal caplet-transmitted gamma-proteobacterial gut symbionts have been characterized in chrysomelid tortoise and reed beetles, these lineages were not detected in the chrysomelids included in these analyses [51, 52].

### Concluding remarks

While insect-microbial symbioses are well established in a few host species, this unprecedented present study sought to apply a phylosymbiosis framework, which suggests that the relationships between hosts and their microbiomes exist due to host evolution as opposed to random chance, to examine eco-evolutionary patterns over ∼400 My of metazoan history and discern how microbiome compositionality has evolved over a multitude of host speciation events. With a reasonably well-defined insect host phylogeny in-hand, it was possible to evaluate phylosymbiosis across major insect orders by measuring the relationship between host phylogenetic dissimilarity and microbiome composition dissimilarity. While mammals, birds, and other terrestrial animals, including insects, have been the subjects of past efforts to explain host-microbiome interactions through a phylosymbiosis lens, this is the first study to include lineages representing this ancestral depth of animal evolution. The pan-Insecta phylosymbiosis r-statistic was driven by large differences between Blattodea and other insect orders rather than by a tractable pattern of microbiome changes across insect host evolution, and sister taxa were often observed to not harbor similar bacterial communities **(Fig. 4).** As such, phylosymbiosis metrics as used in this study may have a ceiling-of-reliability given the phylogenetic range of included host species. Despite this instability, inter- and intra-generational bacterial transmission modalities and host behaviors (i.e. sociality, gregarious domiciling, etc.) were frequently associated with the consistent presence and prevalence of core phylotypes that defined host species within insect orders and accounted for detected phylosymbiotic signal. Within insect lineages where transmission modalities are unknown or likely absent (e.g. Diptera and Coleoptera), there was a reliable lack of phylosymbiotic signal, which may be expected due to the lack of clear inheritance mechanism between generations and lack of host bacterial filtering. However, the sum of this study’s results indicate that caution may be warranted when attempting to apply phylosymbiotic theory across host species with large variances in (1) microbiome diversity and (2) ability to share bacteria from generation to generation due to the potential to uncover unreliable results.

## DATA AVAILABILITY

All data and analysis code used for this study are available on GitHub https://github.com/kjbuffin/Pan-Insect-Phylosymbiosis. Raw sequence data are available on the NCBI sequence read archive and can be accessed using the accession numbers found in **Supplemental Materials**.

## AUTHOR CONTRIBUTIONS

ZLS and KJB designed research, performed research, analyzed data and wrote the paper.

## REFERENCES

1. Ley RE, et al. Evolution of Mammals and Their Gut Microbes. Science 2008;320:1647–1651. 10.1126/science.1155725

2. Brucker RM, Bordenstein SR. The Roles of Host Evolutionary Reltionships (Genus: Nasonia) and Development in Structuring Microbial Communities. Evol 2012;66:349–362. 10.1111/j.1558-5646.2011.01454.x

3. Brooks AW et al. Phylosymbiosis: Relationships and Functional Effects of Microbial Communities across Host Evolutionary History. PLOS Biology 2016;14:e2000225. 10.1371/journal.pbio.2000225

4. Shapira M. Gut Microbiotas and Host Evolution: Scaling Up Symbiosis. Trends in Ecology & Evolution 2016;31:539–549. 10.1016/j.tree.2016.03.006

5. Henry LP et al. The microbiome extends host evolutionary potential. Nat Commun 2021;12:5141. 10.1038/s41467-021-25315-x

6. Lim SJ, Bordenstein SR. An introduction to phylosymbiosis. Proceedings of the Royal Society B: Biological Sciences 2020;287:20192900. 10.1098/rspb.2019.2900

7. Song SJ et al. Comparative Analyses of Vertebrate Gut Microbiomes Reveal Convergence between Birds and Bats. mBio 2020;11:e02901–19. 10.1128/mBio.02901-19

8. Grond K et al. No evidence for phylosymbiosis in western chipmunk species. FEMS Microbiol Ecol 2020;96:fiz182. 10.1093/femsec/fiz182

9. Brown BRP et al. Host phylogeny and functional traits differentiate gut microbiomes in a diverse natural community of small mammals. Molecular Ecology 2023;32:2320–2334. 10.1111/mec.16874

10. Syoc EPV et al. Gut fungal profiles reveal phylosymbiosis and codiversification across humans and nonhuman primates. PLOS Biology 2025;23:e3003390. 10.1371/journal.pbio.3003390

11. Trevelline BK et al. A bird’s-eye view of phylosymbiosis: weak signatures of phylosymbiosis among all 15 species of cranes. Proc Biol Sci 2020;287:20192988. 10.1098/rspb.2019.2988

12. Tinker KA, Ottesen EA. Phylosymbiosis across Deeply Diverging Lineages of Omnivorous Cockroaches (Order Blattodea). Applied and Environmental Microbiology 2020;86:e02513–19. 10.1128/AEM.02513-19

13. Jackson R et al. Evidence of phylosymbiosis in Formica ants. Front Microbiol 2023;14. 10.3389/fmicb.2023.1044286

14. O’Brien PA et al. Diverse coral reef invertebrates exhibit patterns of phylosymbiosis. ISME J 2020;14:2211–2222. 10.1038/s41396-020-0671-x

15. Perez-Lamarque B et al. Limited Evidence for Microbial Transmission in the Phylosymbiosis between Hawaiian Spiders and Their Microbiota. mSystems 2022;7:e01104–21. 10.1128/msystems.01104-21

16. Lin L-Q, Tembrock LR, Wang L. Prevalence and underlying mechanisms of phylosymbiosis in land plants. J Plant Ecol 2024;17:rtae051. 10.1093/jpe/rtae051

17. Graham NJ et al. Evidence of Phylosymbiosis in the Microbiome of Conifer Roots. Phytobiomes Journal 2025;9:541–557. 10.1094/PBIOMES-03-25-0022-R

18. Mazel F et al. Is Host Filtering the Main Driver of Phylosymbiosis across the Tree of Life? mSystems 2018;3:10.1128/msystems.00097-18. 10.1128/msystems.00097-18

19. Mallott EK. Disentangling the mechanisms underlying phylosymbiosis in mammals. Molecular Ecology 2024;33:e17193. 10.1111/mec.17193

20. Muegge BD et al. Diet drives convergence in gut microbiome functions across mammalian phylogeny and within humans. Science 2011;332:970–974. 10.1126/science.1198719

21. Yun J-H et al. Insect Gut Bacterial Diversity Determined by Environmental Habitat, Diet, Developmental Stage, and Phylogeny of Host. Appl Environ Microbiol 2014;80:5254–5264. 10.1128/AEM.01226-14

22. Engel P, Moran NA. The gut microbiota of insects – diversity in structure and function. FEMS Microbiology Reviews 2013;37:699–735. 10.1111/1574-6976.12025

23. Martinson VG et al. A simple and distinctive microbiota associated with honey bees and bumble bees. Molecular Ecology 2011;20:619–628. 10.1111/j.1365-294X.2010.04959.x

24. McLean AHC et al. Host relatedness influences the composition of aphid microbiomes. Environ Microbiol Rep 2019;11:808–816. 10.1111/1758-2229.12795

25. Tinker KA, Ottesen EA. The Core Gut Microbiome of the American Cockroach, Periplaneta americana, Is Stable and Resilient to Dietary Shifts. Applied and Environmental Microbiology 2016;82:6603–6610. 10.1128/AEM.01837-16

26. Parker ES, Newton ILG, Moczek AP. (My Microbiome) Would Walk 10,000 miles: Maintenance and Turnover of Microbial Communities in Introduced Dung Beetles. Microb Ecol 2020;80:435–446. 10.1007/s00248-020-01514-9

27. Moran NA, Baumann P. Bacterial endosymbionts in animals. Current Opinion in Microbiology 2000;3:270–275. 10.1016/S1369-5274(00)00088-6

28. Gil R et al. The genome sequence of Blochmannia floridanus: Comparative analysis of reduced genomes. Proceedings of the National Academy of Sciences 2003;100:9388–9393. 10.1073/pnas.1533499100

29. Aksoy S. Wigglesworthia gen. nov. and Wigglesworthia glossinidia sp. nov., Taxa Consisting of the Mycetocyte-Associated, Primary Endosymbionts of Tsetse Flies. International Journal of Systematic and Evolutionary Microbiology 1995;45:848–851. 10.1099/00207713-45-4-848

30. Moran NA, McCutcheon JP, Nakabachi A. Genomics and Evolution of Heritable Bacterial Symbionts. Annual Review of Genetics 2008;42:165–190. 10.1146/annurev.genet.41.110306.130119

31. Bansal R, Michel AP, Sabree ZL. The crypt-dwelling primary bacterial symbiont of the polyphagous pentatomid pest Halyomorpha halys (Hemiptera: Pentatomidae). Environ Entomol 2014;43:617– 625. 10.1603/EN13341

32. Nalepa CA, Bignell DE, Bandi C. Detritivory, coprophagy, and the evolution of digestive mutualisms in Dictyoptera. Insectes soc 2001;48:194–201. 10.1007/PL00001767

33. Itoh H et al. Soil pH as an external filter shaping stink bug–Burkholderia gut symbiosis. Microbiome 2026;14:129. 10.1186/s40168-026-02402-z

34. Kaltenpoth M et al. Origin and function of beneficial bacterial symbioses in insects. Nat Rev Microbiol 2025;1–17. 10.1038/s41579-025-01164-z

35. Bolyen E et al. Reproducible, interactive, scalable and extensible microbiome data science using QIIME 2. Nat Biotechnol 2019;37:852–857. 10.1038/s41587-019-0209-9

36. Martin M. Cutadapt removes adapter sequences from high-throughput sequencing reads. EMBnet.journal 2011;17:10–12. 10.14806/ej.17.1.200

37. Callahan BJ et al. DADA2: High-resolution sample inference from Illumina amplicon data. Nat Methods 2016;13:581–583. 10.1038/nmeth.3869

38. Chuvochina M, et al. SILVA in 2026: a global core biodata resource for rRNA within the DSMZ digital diversity. Nucleic Acids Res 2026;54:D334–D341. 10.1093/nar/gkaf1247

39. Oksanen J et al. vegan: Community Ecology Package. 2026. 2026.

40. Kumar S et al. TimeTree 5: An Expanded Resource for Species Divergence Times. Mol Biol Evol 2022;39:msac174. 10.1093/molbev/msac174

41. Bourguignon T et al. Transoceanic Dispersal and Plate Tectonics Shaped Global Cockroach Distributions: Evidence from Mitochondrial Phylogenomics. Mol Biol Evol 2018;35:970–983. 10.1093/molbev/msy013

42. Bell WJ, Roth LM, Nalepa CA. Cockroaches: Ecology, Behavior, and Natural History. JHU Press, 2007.

43. Arora J et al. Evidence of cospeciation between termites and their gut bacteria on a geological time scale. Proc Biol Sci; 290:20230619. 10.1098/rspb.2023.0619

44. Piquer-Esteban S et al. Blattella germanica Selects Microbiota Taxa from Feces and Environmental Inputs. Insects 2026;17:615. 10.3390/insects17060615

45. Jahnes BC, Herrmann M, Sabree ZL. Conspecific coprophagy stimulates normal development in a germ-free model invertebrate. PeerJ 2019;7:e6914. 10.7717/peerj.6914

46. Chouvenc T et al. Termite evolution: mutualistic associations, key innovations, and the rise of Termitidae. Cell Mol Life Sci 2021;78:2749–2769. 10.1007/s00018-020-03728-z

47. Bogri A et al. Transmission of antimicrobial resistance in the gut microbiome of gregarious cockroaches: the importance of interaction between antibiotic exposed and non-exposed populations. mSystems 2023;9:e01018–23. 10.1128/msystems.01018-23

48. Wada-Katsumata A et al. Gut bacteria mediate aggregation in the German cockroach. Proceedings of the National Academy of Sciences 2015;112:15678–15683. 10.1073/pnas.1504031112

49. Prasad A et al. Evolution of gut microbiota across honeybee species revealed by comparative metagenomics. Nat Commun 2025;16:9069. 10.1038/s41467-025-64115-5

50. Kohl KD. Ecological and evolutionary mechanisms underlying patterns of phylosymbiosis in host-associated microbial communities. Philos Trans R Soc Lond B Biol Sci 2020;375:20190251. 10.1098/rstb.2019.0251

51. Reis F et al. Bacterial symbionts support larval sap feeding and adult folivory in (semi-)aquatic reed beetles. Nat Commun 2020;11:2964. 10.1038/s41467-020-16687-7

52. Salem H et al. Drastic Genome Reduction in an Herbivore’s Pectinolytic Symbiont. Cell 2017;171:1520–1531.e13. 10.1016/j.cell.2017.10.029

