## Supplemental text for "Evolutionary relatedness is a partially reliable predictor of host-associated microbiome composition"

**This PDF file includes:**

Supplemental text

Figures S1A-E to S3

Legends for Datasets S1 to S5

**Other supporting materials for this manuscript include the following**

Datasets S1 to S5

**Supplemental Text**

**Supplemental Methods**

*Meta-analysis approach*

A comprehensive search of publicly available sequence data was conducted to construct a dataset comprising the widest range of well-characterized insect bacterial communities possible. Specifically, searches were done on the National Center for Biotechnology Information (NCBI) Sequence Read Archive, to identify BioProjects potentially characterizing the associated internal microbiota of any insect host. Once a comprehensive list of potential projects was generated, manual curation of those BioProjects was done. To survey a wide phylogenetic range of host insects, we limited the included projects to those using Illumina sequencing platforms but kept projects which targeted any 16S gene segment, which we would address through our use of genus-level taxonomic assignment. We heavily utilized the sample associated metadata available for each project from NCBI, as well as any publication referencing each project, to build sample-associated metadata **(Datatset S4)** and remove specific samples and entire projects not relevant to this study. This included samples taken from non-internal tissue, or environmental samples, in addition to samples which included experimental antibiotic treatment. Entire projects were not included because of the inability to differentiate samples across treatments, insect life stages, or other isolation sources. A full list of BioProjects included in the final dataset is available **(Dataset S3)**. We acknowledge that this methodology provides little control over various aspects of ribosomal amplicon sequencing known to affect the resulting characterizations of bacterial communities; including, tissue storage procedures, DNA extraction protocols, and sequencing chemistry. However, we believe important to addressing our focal questions was the ability to include a wide phylogenetic range of host insects. Additionally, as discussed below, we believe both our conservative treatment of the sequencing data and the statistical analyses used attempt to account for expected variance between samples and projects.

*Amplicon Sequence Processing*

Sequencing data were obtained from the NCBI Sequence Read Archive using the SRA Toolkit (version 3.0.0) in FASTQ format. Both paired-end and single-end sequencing strategy BioProjects were included. For datasets containing paired-end reads, only forward reads were retained for downstream analysis for better comparability across all datasets. Each project was independently imported and analyzed within the QIIME 2 framework (version 2024.10) (1). To find primer sequences across reads from our set of diverse projects, a library of common 16S primer sequences as well as those explicitly cited by publications referencing our target projects was constructed. The Cutadapt plugin (2) was then used for trimming. Following primer removal, we applied a conservative quality filtering using QIIME 2 quality-filter q-score plugin with a minimum quality threshold of 15 and a sliding window of 4 base pairs to remove low-quality regions. Amplicon sequence variants (ASVs) were identified using the DADA2 (3) plugin. No additional read trimming was applied at the DADA2 denoising step, and a minimum fold parent-over-abundance threshold of 4 was used to distinguish true biological variants from sequencing errors. Taxonomic classification of ASVs was performed using a pre-trained Bayes classifier based on the SILVA 138 database (99% similarity, full-length sequences, 2024 release) (4) using the classify-sklearn plugin. To ensure retention of only bacterial sequences, multiple filtering steps were applied sequentially. ASVs lacking phylum-level taxonomic assignment were removed, in addition to reads assigned as eukaryotic, chloroplast, or mitochondrial. After read filtering, ASVs were clustered at 100% sequence identity. The clustered reads were then filtered again to remove low-abundance ASVs using a minimum frequency threshold of 11 reads across all samples, removing potentially spurious sequences. Finally, samples with fewer than 10,000 total reads after all filtering steps were excluded to ensure adequate sequencing depth for downstream community analyses. Following individual processing of each BioProject, ASV tables, representative sequences, and taxonomic classifications were merged in QIIME 2. The merged feature table, which included all samples, was collapsed to genus level phylotypes to better allow for inter-project comparisons while maintaining sufficient taxonomic resolution for ecological interpretation. Genus-level phylotypes were the taxonomic units used in this meta-analysis to harmonize data drawn from unrelated studies at the cost of resolution possible with species-level phylotypes or ASVs.

*Host phylogeny and metadata*

We generated a phylogeny of host insects relevant to this dataset using TimeTree (version 5.0) (5). Briefly, we submitted a list of target insect hosts. Most of them were able to be placed onto the phylogeny with the exact matching species, while some hosts needed substitution with closest available species, or were ultimatley excluded entirely based on no closely related entry available in the databse. The host phylogeny was forced to be ultramertic using the force.ultrametric function from phytools 2.0 (version 2.4-4) (6) to ensure appropriate temporal scaling for phylogenetic comparative analyses. Host dietary metadata **(SI – Dataset S4)** was determined based on a priori knowledge of well-documented insects in addition to internet searches for less commonly characterized species. Best approximations were made based on available information for all species. Diets were divided into more broad and more finely defined groups. We classified insects as Carnivorous, if we believed their diet best fit into the narrower categories of hematophagous (both sex are blood feeding), or insectivorous (consumes insects. Insects were classified as Omnivorous if their diet was grouped as coprophagic (dung beetles), female sex only hematophagus insects, or as omnivorous generalists who generally feed on both plant material and insects. Finally, Herbivorous insects included those grouped as Mycophagous (consuming fungi), phloem-feeders, phytophagus, pollen/nectar feeders, and xylophagus (wood feeding).

*Statistical and Phylogenetic Analysis*

Statistical analyses and data visualizations were performed in R (version 4.4.1) (7). The merged genus-level ASV table, taxonomic classifications, and sample metadata were imported into R using phyloseq (version 1.48.0) (8). The per-sample metadata, including source BioProject, insect host, and specimen information such as life stage, tissue, sex, and habitat was imported from a manually curated csv file **(Datatset S4)**. Per-host metadata including dietary classifications were also imported from a manually curated csv file (**Dataset S5)**. The host phylogenetic tree was imported in Newick format**.** Multiple filtering steps were applied to the phyloseq object to generate a dataset for analysis. First, non-adult life stages (larvae, nymphs, pupae) were excluded to control for developmental variation in microbiome composition, retaining only adult specimens for analysis. Second, host species with insufficient sample sizes, in our case defined as those with less than 10 samples, were removed to allow for robust intra-host comparisons.

*Identification of core bacterial phylotypes*

Core bacterial phylotypes were hypothesized to be those closely associated with host life history and speciation. Identification of the core microbiome (i.e. the community of core phylotypes) across samples and across different projects accounts for the expected abundance variance associated with destructive DNA extraction protocols. To identify core phylotypes across insect hosts, we employed a bootstrap-based methodology. This approach accounts for unequal sample sizes across host species and provides statistical confidence in our determination of core and non-core phylotypes, and we developed this procedure to account for inter-sample variations within each host.

To summarize, genus-level phyloseq objects were transformed to binary presence/absence data (any non-zero abundance = 1, zero abundance = 0). Classifying core phylotypes by frequency of detection, rather than by relative abundance alone, minimized temporary and/or stochastic conditions (i.e. feeding status) that may skew the relative abundance of a gut bacterial lineage at the time of sampling. For each host species, we first identified if different levels existed across five sample metadata factors: tissue, sex, habitat, life stage, and source BioProject. If any of those factors had a level reaching 10 samples, that subset also had this procedure performed on it. For a given host’s entire sample set, and any applicable factor level sample sets, 1,000 bootstrap iterations were performed. In each iteration, samples were randomly drawn with replacement from the available samples for that host species. In each bootstrap iteration, the proportion of samples containing each bacterial phylotype was calculated. Based on those iterations, a 95% confidence interval of prevalence was determined from the bootstrap distribution for each bacterial phylotype. We utilized a threshold of 80% to call core phylotypes per host. In other words, a bacterial phylotype was considered a core phylotype within a given host species if 0.8 fell within its bootstrapped confidence interval.

*Incidence of core bacteria across host species*

To assess whether core bacteria are shared across a range of host species, we counted the total number of both host families and host species each core phylotype was found in. We additionally determined the average relative abundance of core phylotypes for each host species by computing the sum relative abundance all core bacteria made up for each host’s sample set. Finally, we examined taxonomic diversity of core phylotypes by determining the number of represented bacterial phyla each host’s core community constitutes, in addition to the diversity of phylotypes comprising those phyla.

*Measuring bacterial community dissimilarity between host species*

To distinguish between consistently host-associated bacterial lineages and transient or environmentally acquired taxa, we partitioned the microbiome into three operational categories based on prevalence. First, our whole community partition included all bacteria within each sample. The core partition included only those bacteria we determined to be core phylotypes (in at least 80% of samples for a given host). While the non-core partition was the inverse. We used all 3 of these partitions to measure dissimilarity between host communities in addition to utilizing both abundance weighted and binary data structuring. Abundance weighted dissimilarity comparisons were made using Aitchison distance, by centerlog ratio transforming relative abundances using the microbiome R package (version 1.26.0) (9). Binary dissimilarity comparisons were made using Jaccard dissimilarity calculated by vegdist (vegan package, version 2.7-1). For all six of these community comparisons, a script was utilized to find the centroid of each host’s samples followed by generating the pairwise matrix of centroid to centroid distances between hosts. Alternatively, we also calculated abundance-weighted dissimilarity by averaging the relative abundances of bacterial phylotypes for all samples in each host individually, followed by determining the Aitchison distance between each of those summarized compositions. However, we found that the centroid method resulted in a stronger phylosymbiosis result, while likely also better accounting for intra-host variances and providing a more comparable approach between abundance weighted and binary methods. To visualize compositional dissimilarity between host species, we took our binary dissimilarity matrices and applied hierarchical clustering before plotting as a dendrogram **(Fig. 4)**.

*Testing for phylosymbiosis*

To test for phylosymbiosis we measured the correlation between host phylogenetic relatedness and microbiome dissimilarity by utilizing Mantel tests. The host phylogenetic distance matrix was calculated as the cophenetic distance matrix from the host phylogenetic tree. This matrix quantifies the phylogenetic distance between each pair of host species as the sum of branch lengths separating them on the tree. Mantel tests were conducted separately for each of the six microbiome community partitions (binary and abundance weighted treatments of the whole, core, and non-core partitions) using the mantel function from vegan (version 2.7-1) with 999 permutations to generate null distributions. Statistical significance was assessed by comparing the observed Mantel r to the null distribution generated by permutations.

*Partial Mantel Tests Controlling for Diet*

To determine whether the observed phylogenetic signal was contributed to by dietary similarity rather than direct host phylogenetic effects, we performed partial Mantel tests that statistically account for the effect of diet group. A categorical diet dissimilarity matrix was constructed from a host dietary trait using Gower distance. Partial Mantel tests were conducted using the mantel.partial function from vegan (version 2.7-1) with 999 permutations. This analysis partials out the variance in microbiome composition able to be explained by the diet group variable, isolating the residual phylogenetic signal. The partial Mantel r represents the correlation between host phylogeny and microbiome composition independent of dietary effects. Results were compared to simple Mantel tests to quantify the proportion of phylosymbiosis attributable to phylogenetically conserved diet versus other phylogenetic factors.

*Evaluation of phylosymbiosis robustness*

To assess the robustness of phylogenetic signal, we wanted to determine if specific host orders disproportionately influenced the measured phylosymbiosis across the entire phylogeny. Additionally, we aimed to determine if a strong phylogenetic signal within any insect order was driving the trend across all hosts. To the first point, we performed a jackknife analysis which took each insect order represented in the dataset and removed all host species belonging to that order, before recalculating phylosymbiosis on the remaining species. This procedure was repeated independently for each microbiome partition (whole, core, and non-core community). To evaluate whether phylosymbiosis observed across all insects also manifested within individual insect orders, we conducted order-specific Mantel tests. For each insect order represented by at least 8 host species, we subset the data to include only species from that order. For each qualifying order, the host phylogenetic distance matrix, microbiome dissimilarity matrices (for all three community partitions), and diet dissimilarity matrix were subset to include only species within that order. Both simple and partial Mantel tests were performed using 999 permutations.

**REFERENCES**

1.	E. Bolyen, *et al.*, Reproducible, interactive, scalable and extensible microbiome data science using QIIME 2. *Nat Biotechnol* **37**, 852–857 (2019).

2.	M. Martin, Cutadapt removes adapter sequences from high-throughput sequencing reads. *EMBnet.journal* **17**, 10–12 (2011).

3.	B. J. Callahan, *et al.*, DADA2: High-resolution sample inference from Illumina amplicon data. *Nat Methods* **13**, 581–583 (2016).

4.	M. Chuvochina, *et al.*, SILVA in 2026: a global core biodata resource for rRNA within the DSMZ digital diversity. *Nucleic Acids Res* **54**, D334–D341 (2026).

5.	S. Kumar, *et al.*, TimeTree 5: An Expanded Resource for Species Divergence Times. *Mol Biol Evol* **39**, msac174 (2022).

6.	L. J. Revell, phytools 2.0: an updated R ecosystem for phylogenetic comparative methods (and other things). *PeerJ* **12**, e16505 (2024).

7.	R Core Team, A Language and Environment for Statistical Computing. (2024). Deposited 2024.

8.	P. J. McMurdie, S. Holmes, phyloseq: An R Package for Reproducible Interactive Analysis and Graphics of Microbiome Census Data. *PLOS ONE* **8**, e61217 (2013).

9.	L. Lahti, S. Shetty, Tools for microbiome analysis in R. (2017). Deposited 2017.

**Supplemental Figures**

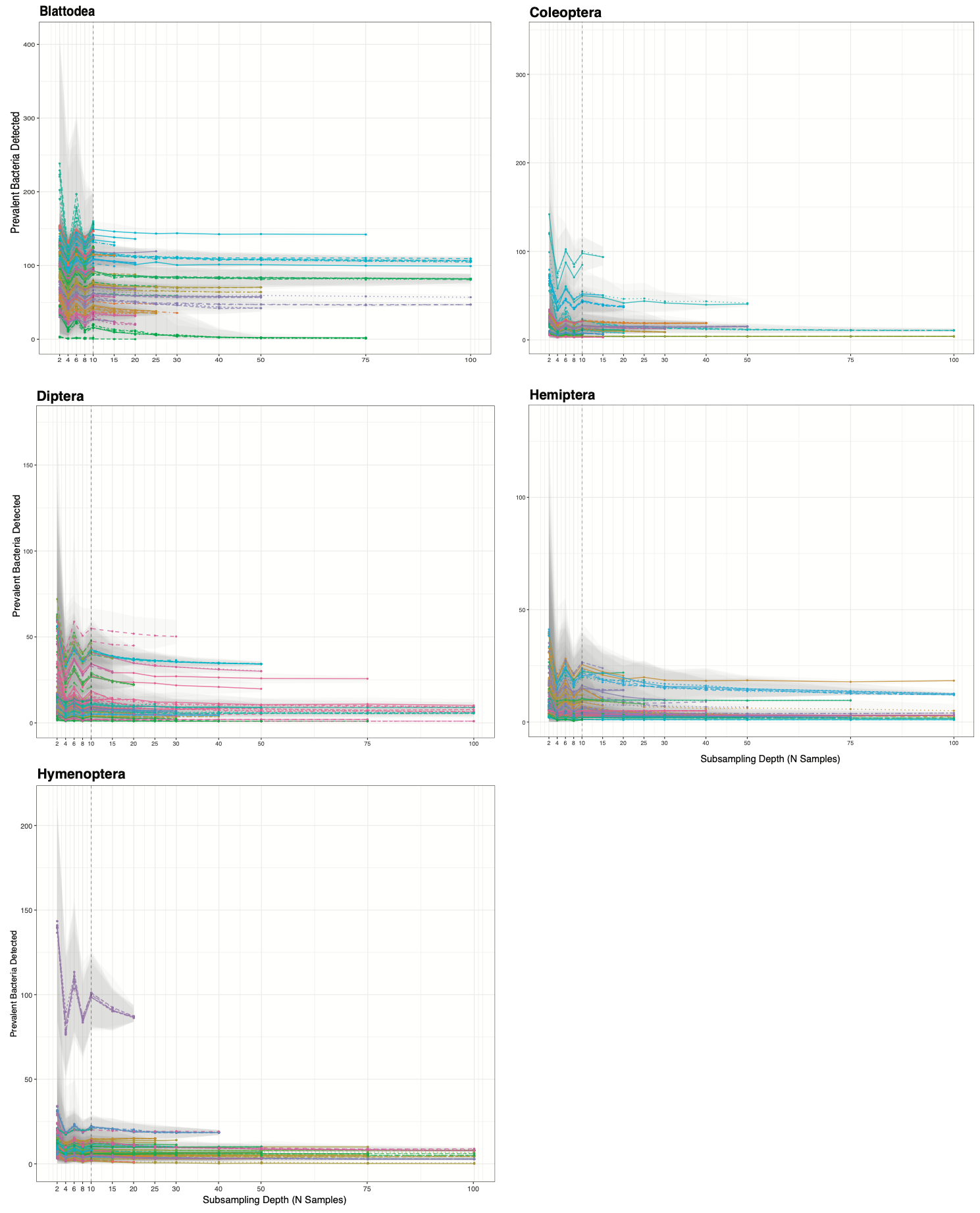

**A**

**B**

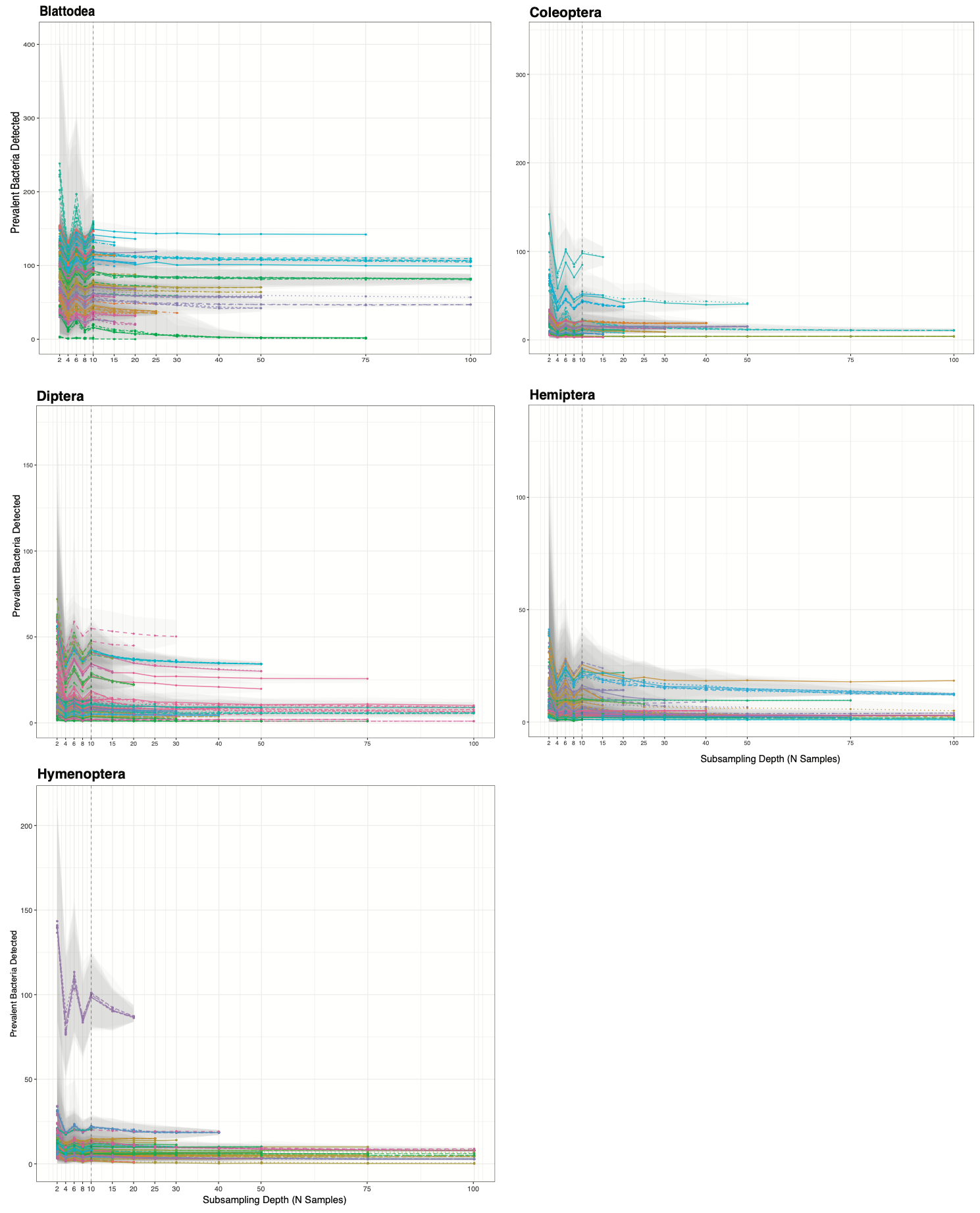

**C**

**D**

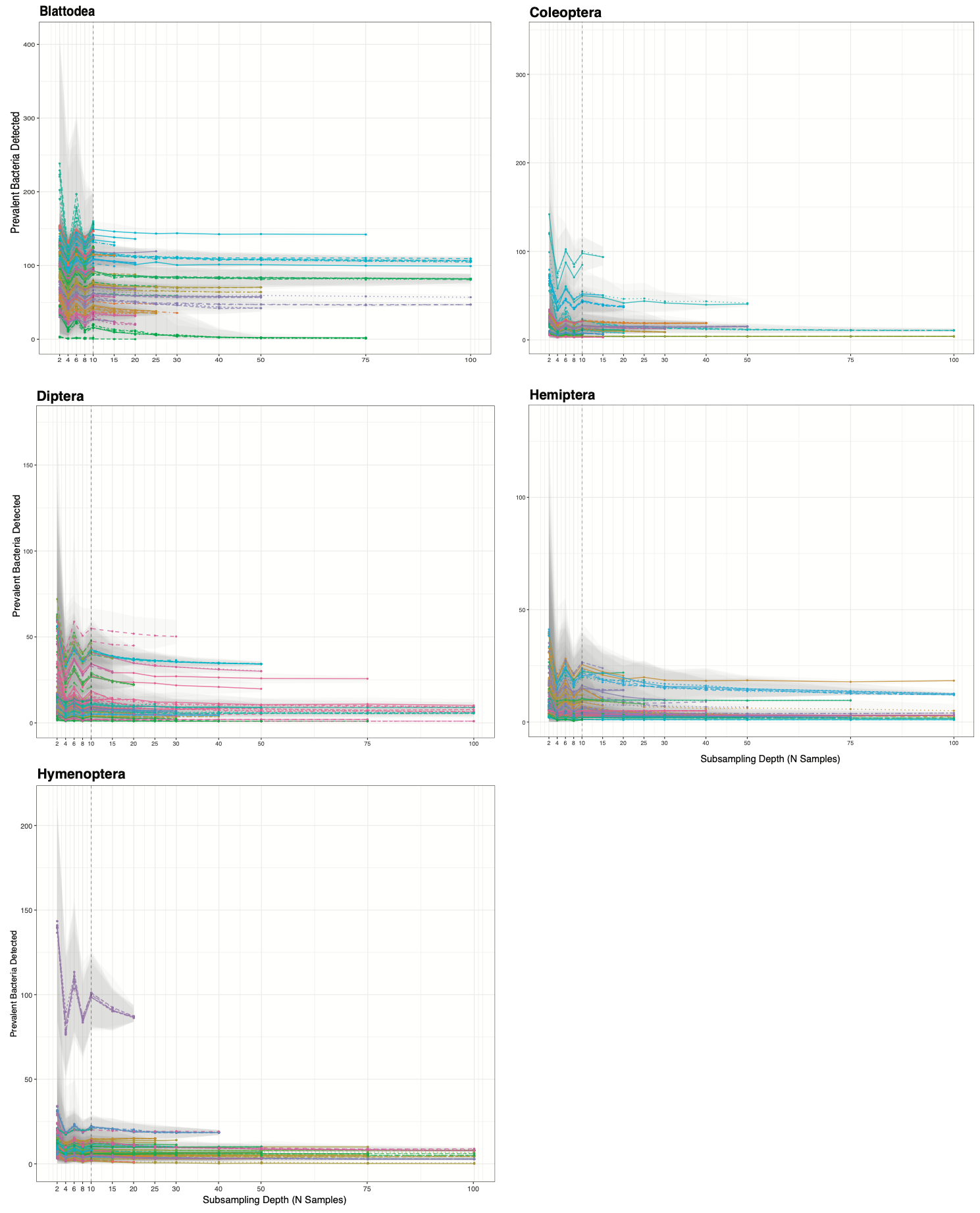

**E**

**Figure S1A-E: Core species accumulation curves with increasing sampling depth**. For each host’s entire sample set or metadata subset, we iteratively ran the prevalence determination at increasing sub-sample counts, each time recording the number of core phylotypes. We observe that around our sample limit of ten, we begin to see leveling of the curve and a decrease in variance across most hosts. The overall trend is that as number of samples is increased during core phylotype determination there is a decrease in number of core taxa called, until a minimum is reached usually between 10 and 20 samples.

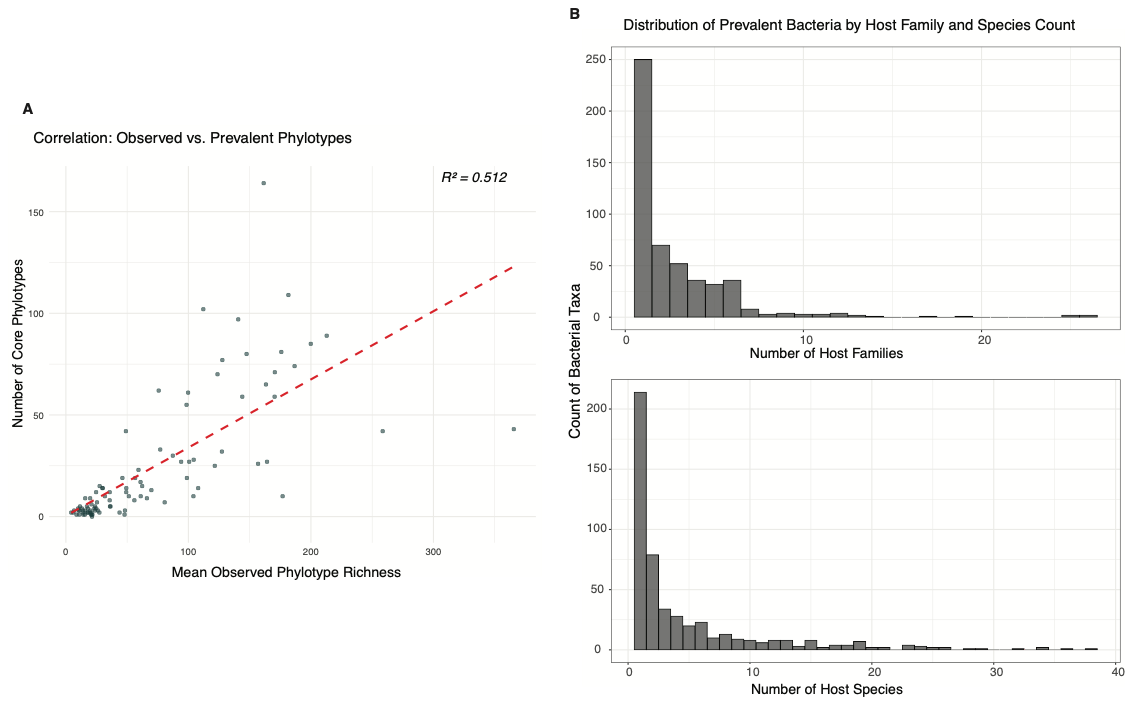

**Figure S2:** **Patterns of core phylotypes distributions across insects**. **A** Correlation demonstrating relationship between average phylotype richness and core phylotypes richness for all insect species. Generally finding that the greater the average whole community richness, the greater the number of core phylotypes. (R2 = 0.513). **B** Distribution of number of host species and host family’s core phylotypes are found in across the insect dataset. Most phylotypes occur in only 1 host species or family, with the distribution being heavily right skewed. Meaning, very few bacterial phylotypes identified as core in our analysis are found in more than one or a couple hosts.

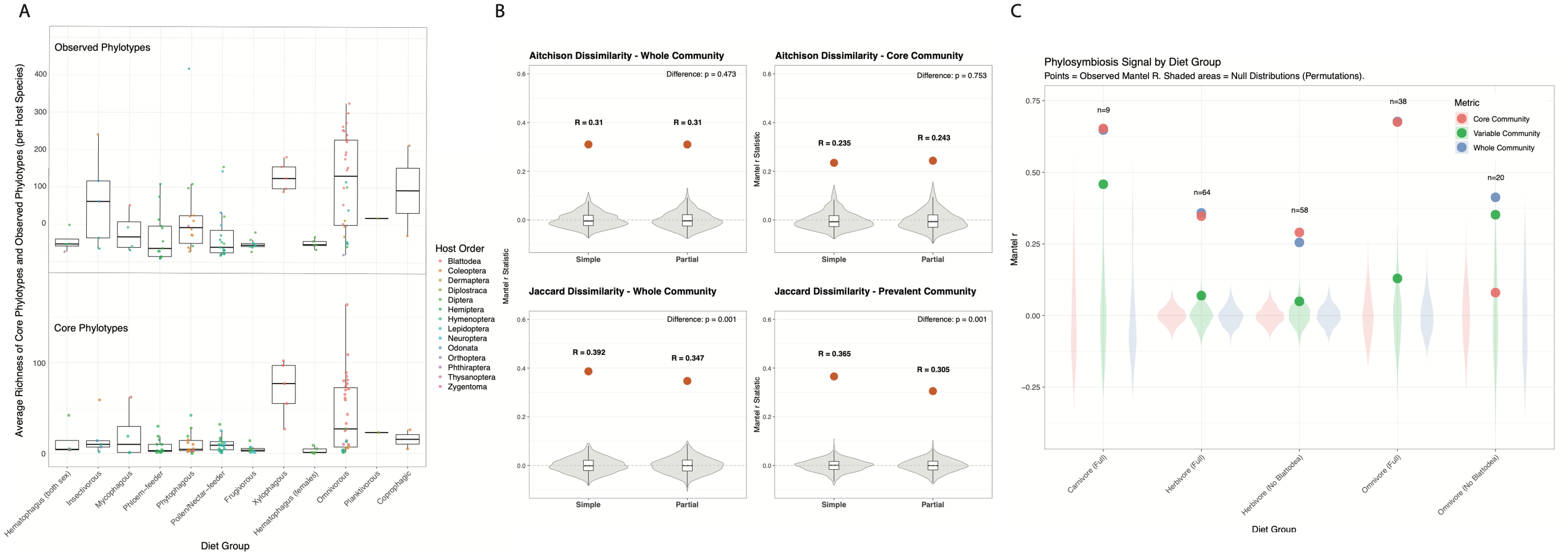

**Figure S3:** **Diet is a driver of bacterial community diversity but plays a limited role in measurement of phylosymbiosis.** **A** For each host species average observed phylotypes and number of core phylotypes were determined. Each insect was assigned one of 13 classifications which generally fall into either carnivorous, herbivorous, or omnivorous and boxplots of diversity are made for each diet group. **B** Partial mantel tests were carried out to evaluate the contribution dietary similarity had on the observed phylosymbiosis. Generally, dietary similarity had a limited effect on the results. **C** Phylosymbiosis was additionally measured within major diet groups (carnivore, omnivore, and herbivore). Because of the known effect Blattodea had on the whole dataset measure, we also removed them from the omnivore and herbivore measurements separately.

**Legends for Datasets S1 – S5**

**S1: Insect microbiome host species details.** All host insects from which microbiome samples were drawn from are listed with their insect order, number of microbiome samples used in these analyses, presumed dietary strategy, average number of observed bacterial genera, number of core bacterial genera, average amplicon reads per sample and number of microbiome sequencing projects used.

**S2: Matrix of core phylotypes per insect host species.** Each genus-level phylotype was determined to be a core phylotype or not in each host species. In this matrix core phylotypes (columns) are represented by a 1 for each insect species (rows). '0' indicates that the phylotype was either not detected or was not a core phylotype in that insect host. Two insect species were found to have no core phylotypes and are not included in this matrix.

**S3: List of NCBI BioProject accession codes for data utilized in this study.** A list of every NCBI BioProject with representative samples in the final analysis dataset. Does not include BioProjects which were evaluated in upstream analysis steps but removed from the study for various reasons.

**S4: Table of metadata associated with each sequence read archive (SRA) accession utilized in this study.** This table includes metadata for the sequencing data of each SRA accession in the final dataset of this study. Metadata was generated (1) directly from the NCBI SRA accession-associated metadata, (2) by interpreting and abstracting the accession-associated metadata, or (3) from information provided in publications associated with the accession.

**S5: Table of diet metadata assigned per host species.** For each host species diet group metadata was assigned based on the known or presumed natural diet classification of each organism (adult life stage specifically). Searches for natural diet information were performed in addition to reliance on prior species knowledge.
